## Supplementary figures and tables for "The structure of the Tad pilus alignment complex reveals a periplasmic conduit for pilus extension"

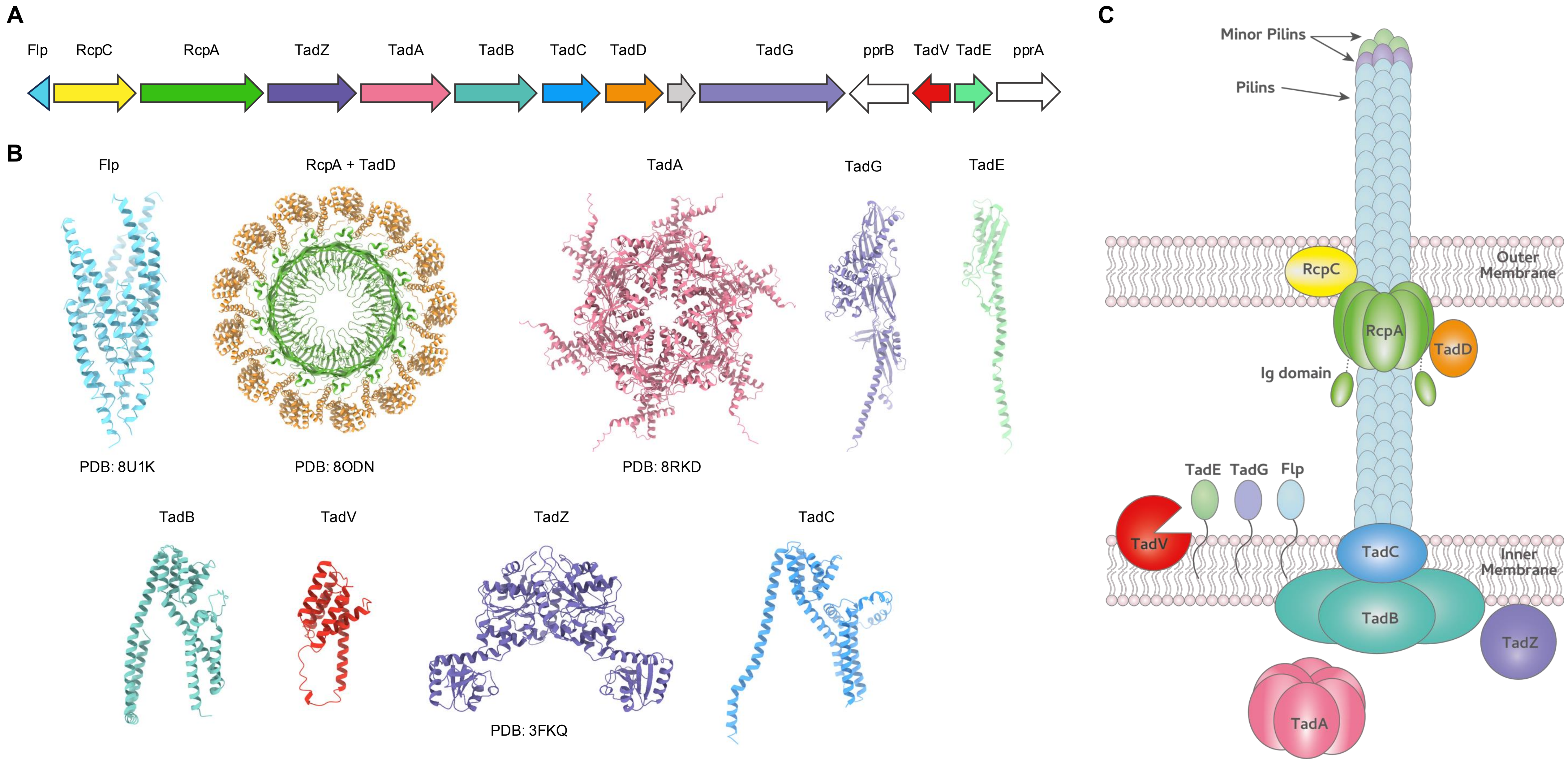


**Extended data figure 1: Organization of the *Pseudomonas aeruginosa* Tad Pilus complex**

**(A)** Tad pilus locus in the Gram-negative bacterium *Pseudomonas aeruginosa*. Genes encoding Tad pilus complex proteins are coloured individually, regulatory genes (*pprA* and *pprB*) are shown in white and uncharacterized genes are indicated in grey. **(B)** Structural models of the corresponding Tad pilus proteins, obtained either from experimental structures (PDB codes indicated) or the AlphaFold database. **(C)** Schematic representation of the Tad pilus complex of *P. aeruginosa*. Individual proteins in **(B)** and **(C)** are coloured as in **(A)**.

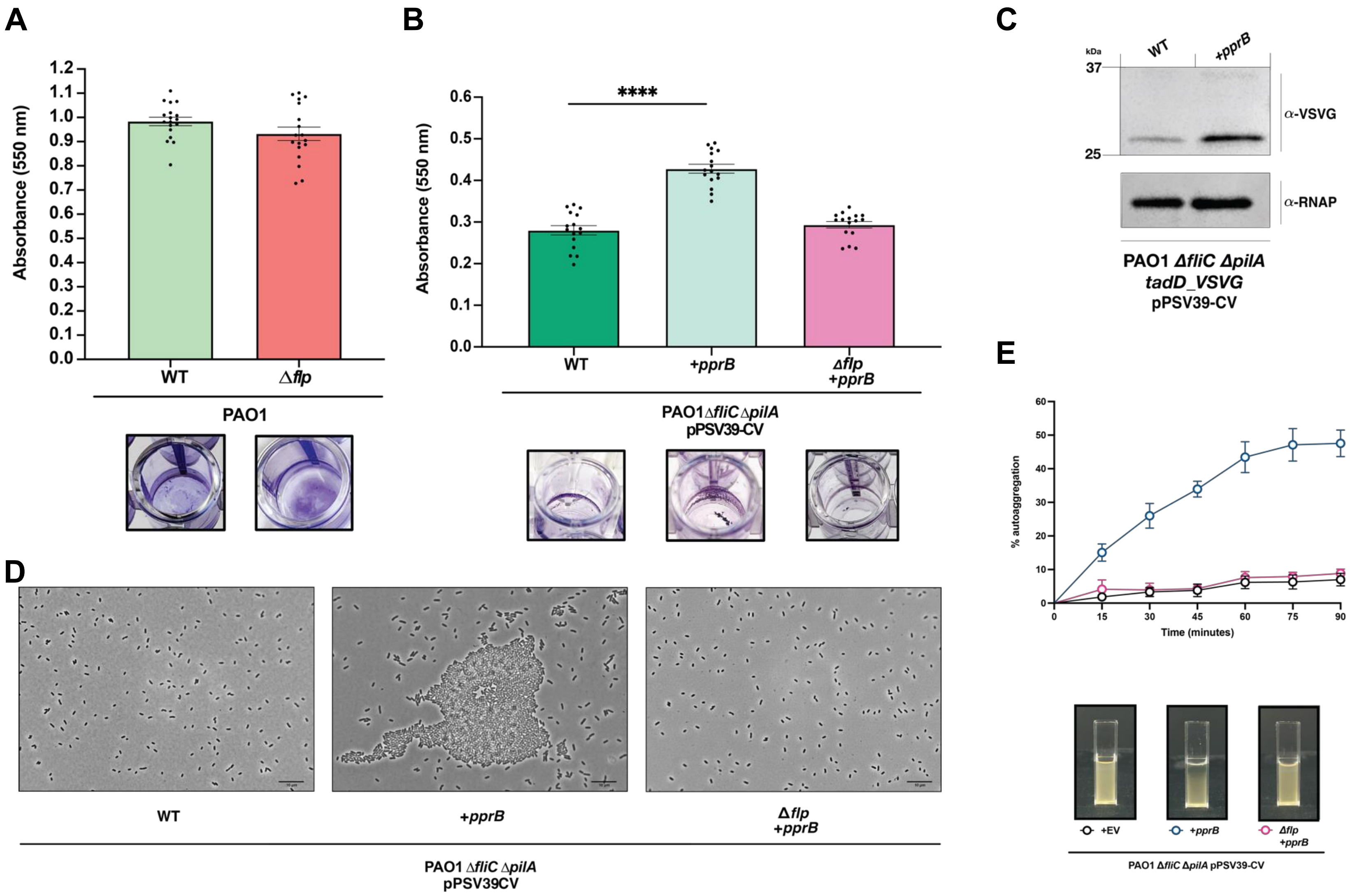


**Extended data figure 2: The Tad pilus induces cell-cell aggregation when over-expressed**

**(A)** Biofilm formation in *P. aeruginosa* WT (left) and Δ*flp* (right) strains. Deletion of the Tad pilus does not affect biofilm formation. **(B)** Overexpression of the Tad pilus transcription activator PprB (centre) leads to a 1.5-fold increase in biofilm formation in comparison to the WT strain (left). This effect is abrogated in a *Δflp* strain (right). **(C)** Overexpression of PprB increases the expression of TadD, demonstrating that this induces the expression of the Tad pilus. **(D)** Phase contrast microscopy of the aforementioned bacterial strains, demonstrating cell-cell aggregation when the Tad pilus is overexpressed. **(E)** Auto-aggregation assay, quantifying the cell-cell aggregation of *P. aeruginosa* strain over-expressing the Tad pilus, dependent on the presence of Flp. Representative views of the cell cultures are shown below.

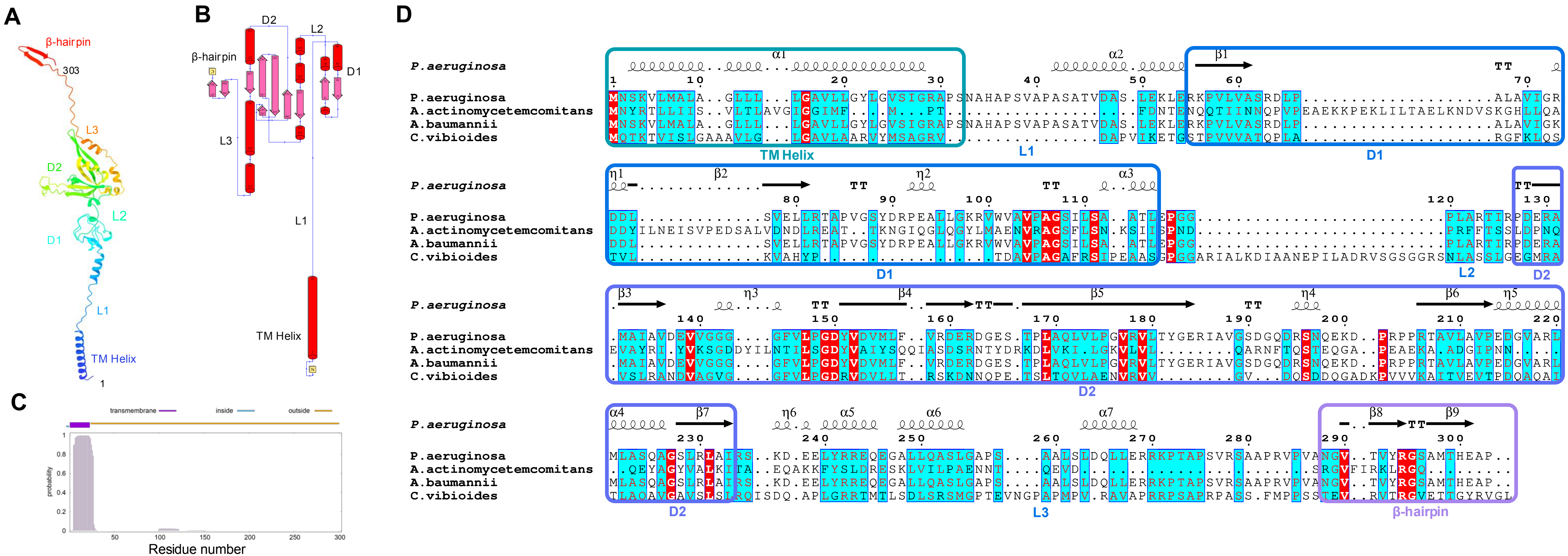


**Extended data figure 3: Structure prediction of RcpC**

**(A)** Cartoon representation, and **(B)** schematic representation, of the RcpC structural model. Its various domains and linkers are indicated. **(C)** Transmembrane prediction analysis of RcpC, indicating that the first ~35 residues have a high probability of forming a transmembrane (TM) helix while the rest of the protein is predicted to be periplasmic. **(D)** Multiple sequence alignment of RcpC showing sequence conservation and similarity across selected species of gram-negative bacteria containing Tad pilus loci. The secondary structure prediction is shown above sequences. Residues highlighted in red are conserved across all species whilst those highlighted in cyan identify residues conserved for most species shown.

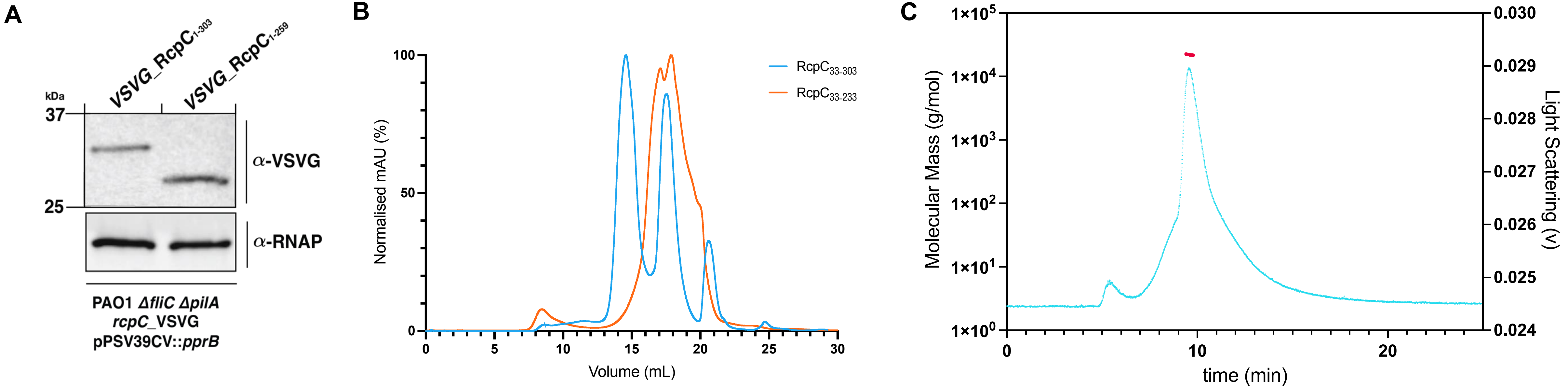


**Extended data figure 4: Characterization of the impact of the deletion of L3 and the β-hairpin**

**(A)** Western blot of $RcpC_{1-303}$ and $RcpC_{1-259}$ complementing the *ΔRcpC P. aeruginosa* strain. α-RNAP is employed as a loading control. **(B)** Gel filtration UV traces for purified $RcpC_{33-303}$ (Cyan) and $RcpC_{33-233}$ (Orange). The peak corresponding to the oligomeric state is not present in the later construct. **(C)** SEC MALS analysis of $RcpC_{33-233}$, with light scattering shown in cyan, and the molecular weight of the corresponding peak shown in red. This corresponds to a molecular weight of ~ 25.8 kDa.

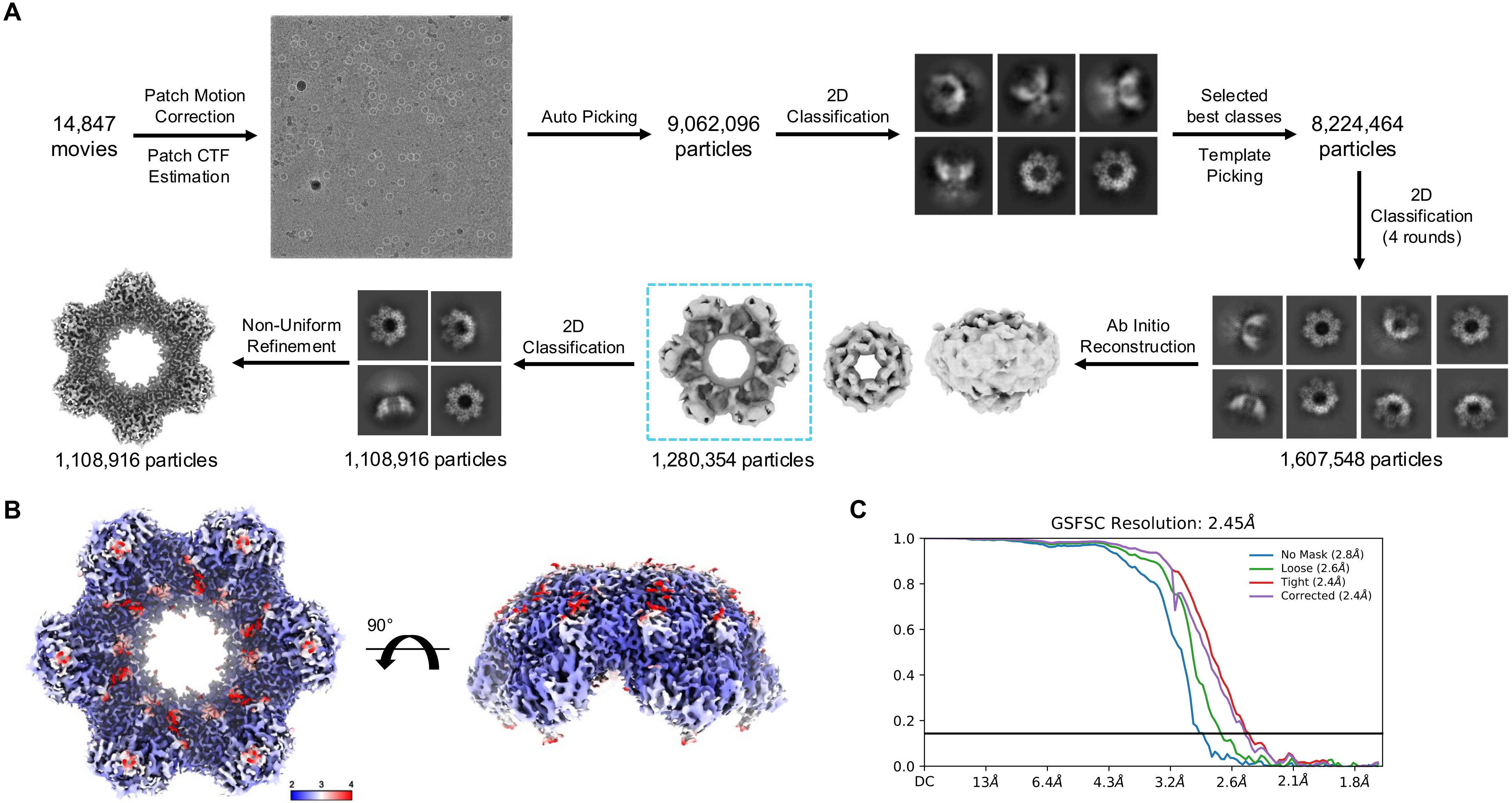


**Extended data figure 5: Cryo-EM processing pipeline for the RcpC dodecamer structure.**
**(A)** Flow chart showing the processing pipeline for the RcpC cryo-EM dataset. The particles were subjected to 4 successive rounds of 2D classification giving the best particles for *ab-initio* reconstruction. The best class form *ab-initio* classification was subjected to further 2D classification from which the particles from the best classes were chosen for Non-Uniform Refinement using C6 symmetry which yielded a 2.5 Å reconstruction. **(B)** Electron potential map of RcpC, coloured by local resolution, showing that most of the map is defined to better than 3 Å. **(C)** FSC curves for the RcpC structure, with C6 symmetry.

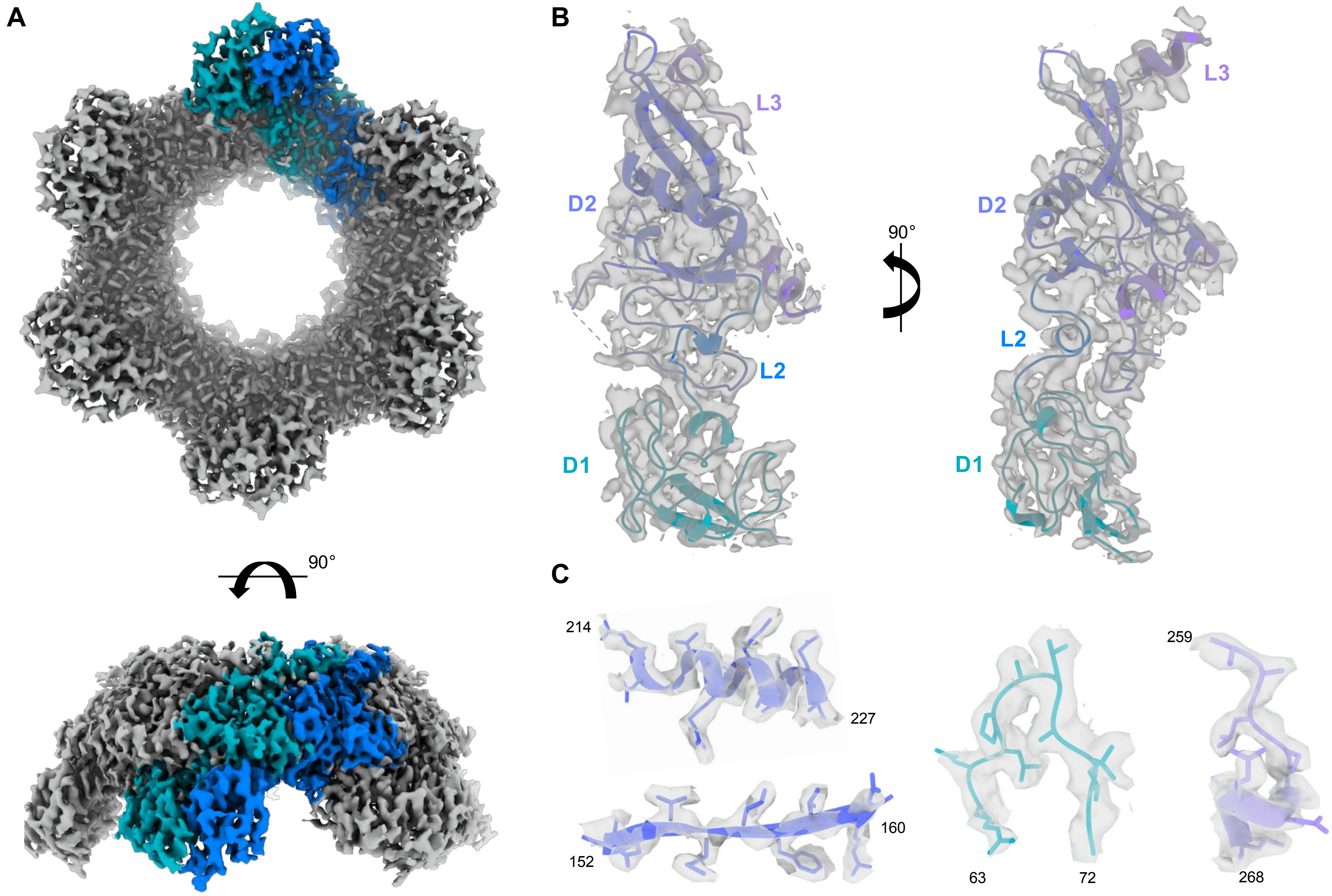


**Extended data figure 6: Cryo-EM map of RcpC.**
**(A)** Electron potential map of RcpC, coloured and segmented for two chains. **(B)** Atomic model of a RcpC molecule, in cartoon representation as coloured in (**A**), with map density shown in transparency. **(C)** Examples of density for four regions of the protein, illustrating that map features correspond to the reported resolution.

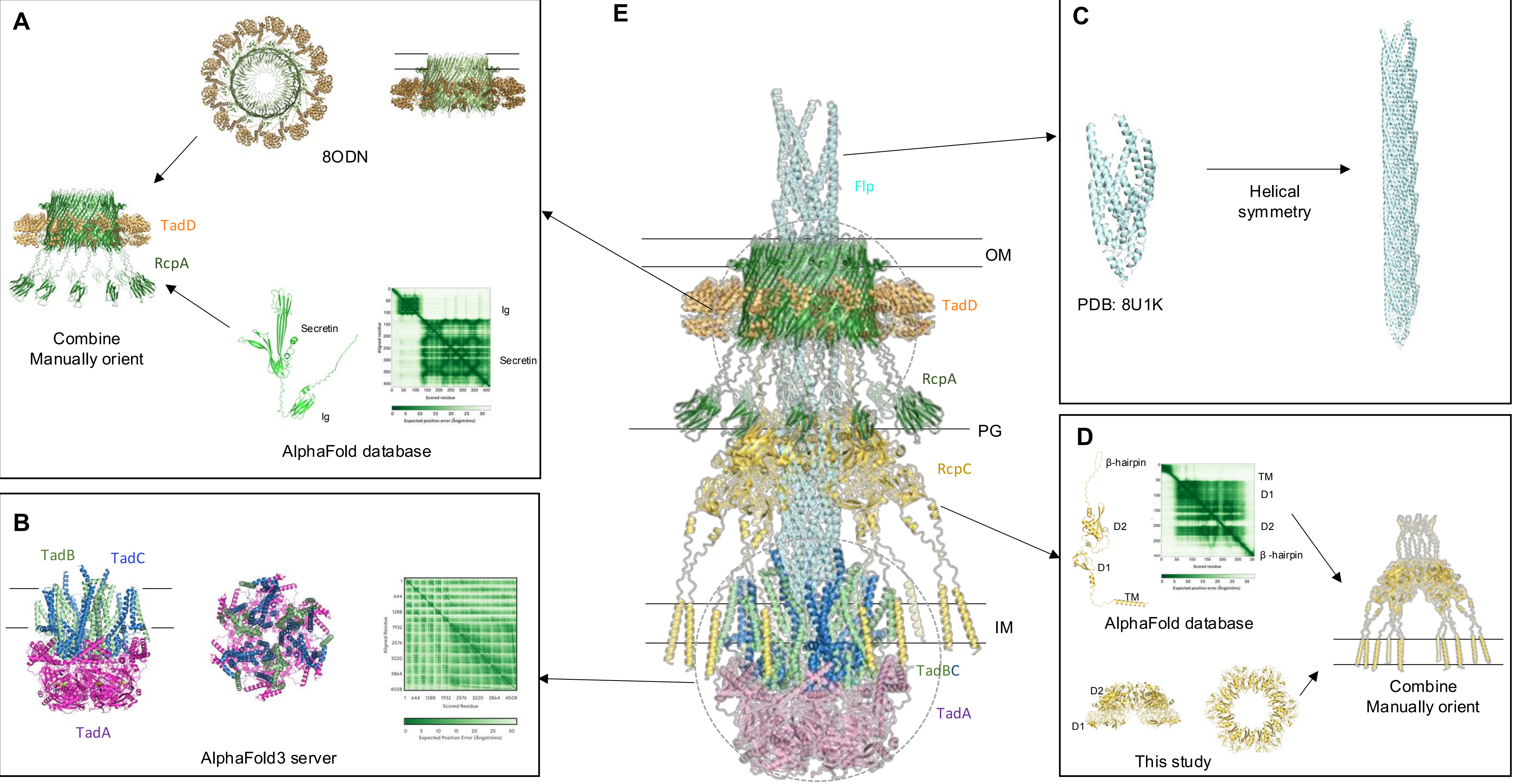


**Extended data figure 7: Modeling the complete Tad pilus assembly.**
**(A)** Composite structure of the full RcpA-TadD complex, obtained by combining the structure of the 13-mer heterodimer complex (PDB ID: 8ODN), and the RcpA full-length model. **(B)** AlphaFold3-generated model of the TadA-TadB-TadC complex. **(C)** Helical structure of the Flp filament (PDB ID: 8U1K). **(D)** Composite structure of the full RcpC complex, obtained by combining the structure of $RcpC_{33-303}$ (this study), and the RcpC full-length model. **(E)** Complete model of the Tad pilus, obtained by combining the aforementioned four models. The localization of of the IM, OM, and peptidoglycan layer (PG) are indicated.

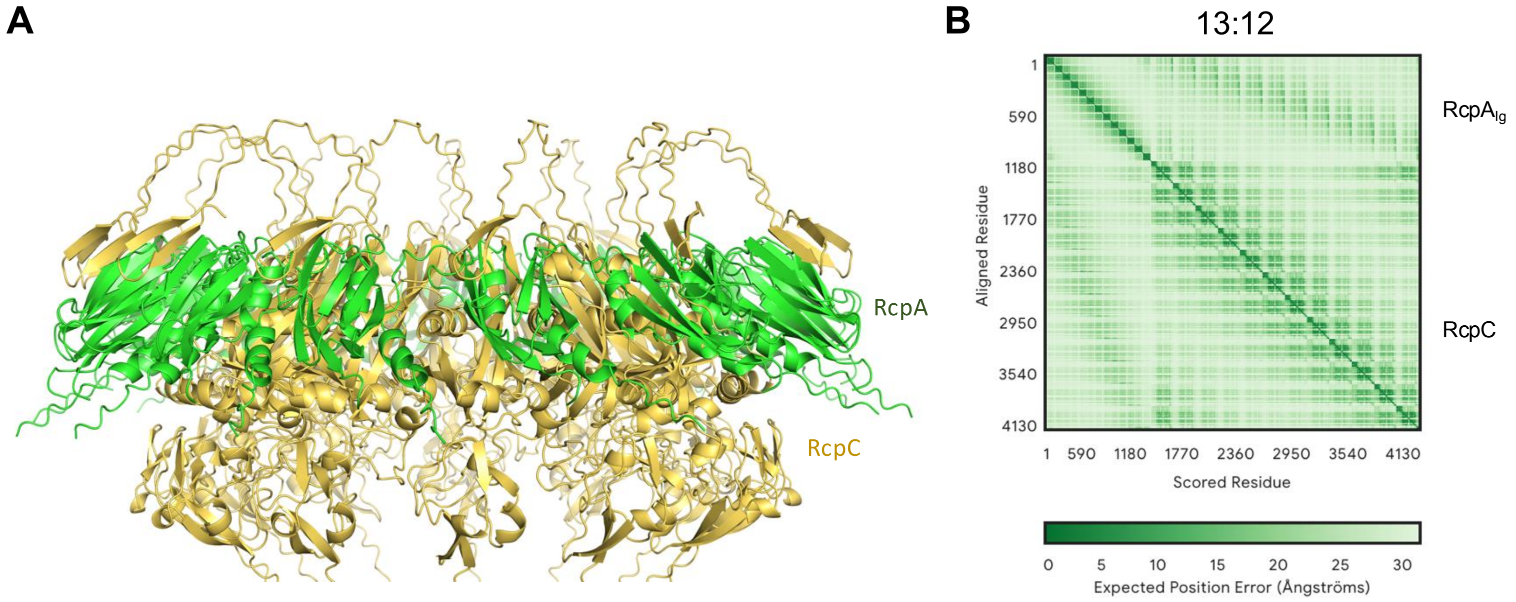


**Extended data figure 8: Modeling of the RcpA-RcpC complex with AlphaFold3**

**(A)** Cartoon representation of the $RcpA_N$-$RcpC_{33\text{-}303}$ complex, modeled with 13:12 stoichiometry, colored in green and yellow, respectively. **(B)** Expected position error for the modeling of this complex, demonstrating that the RcpA-RcpC interface is of high confidence.

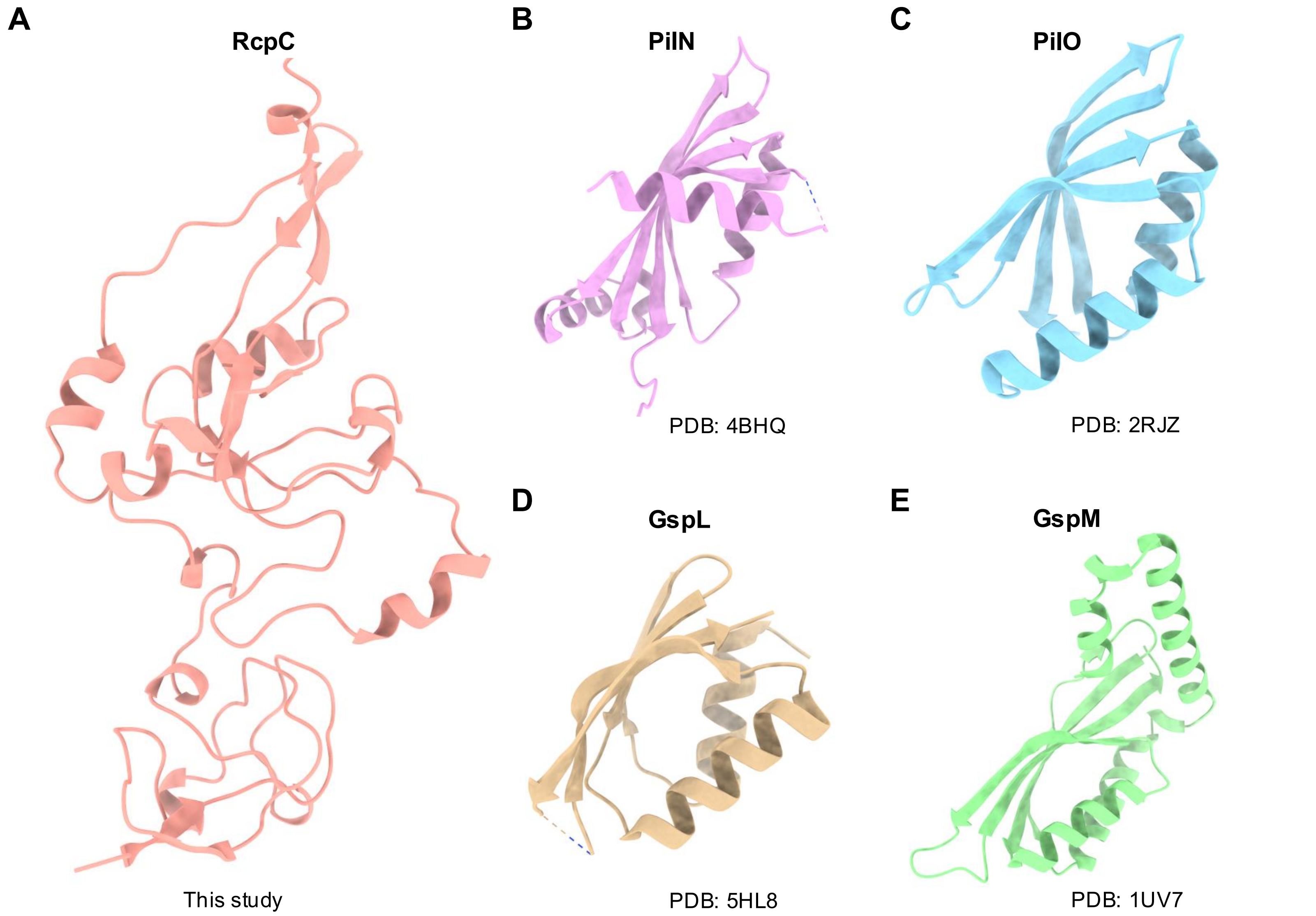


**Extended data figure 9: Comparison of RcpC to other T4P and T2SS alignment complex proteins**
**(A)** Structure of Tad pilus alignment complex protein, RcpC. **(B,C)** Structures of T4P alignment complex proteins PilN (4BHQ) and PilO (2RJZ), respectively. **(D,E)** Structures os T2SS alignment complex proteins GspL (5HL8) and GspM (1UV7), respectively. All structures shown in cartoon representation.

| Data collection | |
|---|---|
| Voltage (kV) | 300 |
| Exposure (e/Å$^2$) | 45 |
| Fractions | 40 |
| Defocus range ($\mu$m) | -0.5 to -2.5 |
| Pixel size (Å/pix) | 0.85 |
| Number of micrographs | 14,847 |
| **Map refinement** | |
| Final particle number | 1,108,916 |
| Resolution (Å) | 2.45 |
| Symmetry | C6 |
| **Structure refinement** | |
| Non-hydrogen atoms | 17,213 |
| Protein residues | 2,272 |
| Protein B-factor | 5.31/ 114.66/ 43.82 |
| Bond length RMSD (Å) | 0.002 |
| Bond angle RMSD (º) | 0.623 |
| MolProbity score | 1.85 |
| Clash score | 4.48 |
| Rotamer outliers (%) | 4.66 |
| Ramachandran favoured (%) | 97.38 |
| Ramachandran allowed (%) | 2.57 |
| Ramachandran disallowed (%) | 0.05 |

**Extended data table 1: Data acquisition and refinement parameters for the RcpC$_{33\text{-}303}$ structure.**

| **Plasmids** | **Description** | **Source** |
|---|---|---|
| pEXG2 | Allelic exchange vector containing both $Gm^R$ *and sacB* | (Rietsch et al., 2005) |
| pPSV39-CV | Derived from pPSV35-CV, containing $Gm^R$ | (Silverman et al., 2013) |
| pEXG2::*ΔfliC* | *fliC* deletion allele in pEXG2 | This study |
| pEXG2::*ΔpilA* | *pilA* deletion allele in pEXG2 | This study |
| pEXG2::*Δflp* | *flp* deletion allele in pEXG2 | This study |
| pEXG2::*tadD_VSVG_Cterm* | Allelic exchange vector including *tadD* with a C-terminal VSV-G chromosomal tag in pEXG2 | This study |
| pEXG2::*rcpC_fulllength_VSVG_G118* | Allelic exchange vector including *rcpC* with a VSV-G tag at residue G118 in pEXG2 | This study |
| pEXG2::*rcpC_VSVG_G118_ ΔCT_A259* | Allelic exchange vector including *rcpC* with a VSV-G tag at residue G118 with the C-terminus truncated from residues 248 to 303 in pEXG2 | This study |
| pPSV39-CV::*pprB* | Expression vector for *pprB* | This study |
| pET28a | Plasmid for protein over-expression | Novagen |
| pET21a | Plasmid for protein over-expression | Novagen |
| pET28a::*rcpC_fulllength* | Over-expression of full-length RcpC, with a C-terminal his-tag | This study |
| pET28a::*rcpC_ΔTM* | Over-expression of $RcpC_{33\text{-}303}$, with a C-terminal his-tag | This study |
| pET28a::*rcpA_NTD* | Over-expression of $RcpA_N$, with a N-terminal his-tag | This study |
| pET21a:: *rcpC_33-303* | Over-expression of $RcpC_{33\text{-}303}$, no tag | This study |
| pET21a:: *rcpC_33-259* | Over-expression of $RcpC_{33\text{-}259}$, no tag | This study |
| pET21a:: *rcpC_33-233* | Over-expression of $RcpC_{33\text{-}233}$, no tag | This study |

**Extended data table 2: List of plasmids used in this study.**

| **Strains** | **Description** | **Source** |
|---|---|---|
| *Pseudomonas aeruginosa* PAO1 | Wild-type *P. aeruginosa* | Goodman et al., 2004 |
| *Pseudomonas aeruginosa* PAO1 *Δflp* | *P. aeruginosa* with deletion of the Tad pilus *flp* gene | This study |
| *Pseudomonas aeruginosa* PAO1 *ΔpilA::ΔfliC* | *P. aeruginosa* with deletion of essential genes for the formation of the T4P and flagellum | This study |
| *Pseudomonas aeruginosa* PAO1 *ΔpilA::ΔfliC::ΔflP* | *P. aeruginosa* with deletion of essential genes for the formation of the T4P and flagellum, and of the Tad pilus *flp* gene | This study |
| *Pseudomonas aeruginosa* PAO1 *ΔpilA::ΔfliC::ΔrcpC* | *P. aeruginosa* with deletion of essential genes for the formation of the T4P and flagellum, and of the Tad pilus *rcpC* gene | This study |
| *Escherichia coli* pLysS | *E. coli* strain for recombinant protein over-expression | Thermo Fisher Scientific |
| *Escherichia coli* DH5α | *E. coli* strain for molecular cloning | Thermo Fisher Scientific |

**Extended data table 3: List of bacterial strains used in this study.**
